# GRATCR: epitope-specific T cell receptor sequence generation with data-efficient pre-trained models

**DOI:** 10.1101/2024.07.21.604503

**Authors:** Zhenghong Zhou, Junwei Chen, Shenggeng Lin, Liang Hong, Dong-Qing Wei, Yi Xiong

## Abstract

T cell receptors (TCRs) play a crucial role in numerous immunotherapies targeting tumor cells. However, their acquisition and optimization present significant challenges, involving laborious and time-consuming wet lab experimental resource. Deep generative model has demonstrated remarkable capabilities in functional protein sequence generation, offering a promising solution for enhancing the acquisition of specific TCR sequences. Here, we propose GRATCR, a framework incorporates two pre-trained modules through a novel “grafting” strategy, to de-novo generate TCR sequences targeting specific epitopes. Experimental results demonstrate that TCRs generated by GRATCR exhibit higher specificity toward desired epitopes and are more biologically functional compared with state-of-the-art model, by using significantly fewer training data. Additionally, the generated sequences display novelty compared to natural sequences, and the interpretability evaluation further confirmed that the model is capable of capturing important binding patterns. GRATCR is freely available at https://github.com/zhzhou23/GRATCR.

## Introduction

T cells, an integral part of the immune system, play a pivotal role in adaptive immunity by recognizing a diverse range of antigens [1, 2]. This recognition is facilitated by the T cell receptor (TCR), a specialized cell surface protein crucial for antigen recognition and T cell activation [3]. Typically, T cell receptor is a heterodimer consisting of alpha and beta chains and the specific recognition of an antigen is primarily achieved through the TCR complementarity-determining region (CDR) [4, 5]. Within CDR, the hypervariable CDR3 loops predominantly interact with the antigenic peptides namely epitopes presented by major histocompatibility complex (MHC) molecules [6–9]. Since TCRs play the important role in the immune process, there are currently a number of promising therapies based on T cell receptors for combatting tumors [10–12]. TCR-T is currently one of the latest and most effective immunotherapies in the treatment of several indications, including large B cell lymphoma, melanoma and other tumors [13, 14]. The treatment utilizes gene editing technology to engineer T cells to express TCRs targeting specific antigens, enabling the recognition and elimination of tumor cells [15, 16]. However, currently TCR-based therapies still encounter significant challenges, and the establishment of a reliable TCR library for targeting specific tumor cells remains a major obstacle [17, 18]. Furthermore, the wet experimental methods require substantial time and financial resources [19, 20]. In the past, numerous deep learning-based models have been applied to TCR-related problems, primarily aiming to elucidate the mechanisms of recognition between TCRs and epitopes. These models focus on determining whether a given TCR and epitope can specifically recognize each other. For instance, ImRex draws inspiration from image classification tasks and utilizes convolutional neural networks (CNN) as the core of the model since CNN is capable of extracting multi-scale features from sequences [21]. TEINet [22], pMTnet [23] and ERGO [24] employ pre-trained models to extract embeddings from input sequences including epitopes and TCRs. Both TEPCAM [25] and TITAN [26] leverage attention mechanisms to capture the interactions between sequences. Although these models demonstrate commendable performance in their respective tasks, they all suffer from poor generalization [27], significantly limiting their practical applications. Moreover, theoretically, the number of TCRs that can be generated by V(D)J recombination far exceeds the number of currently available [28]. Consequently, these models are insufficient for exploring a larger TCR space to expand the current TCR library.

The rapid advancement of generative models and the success of certain biological sequence designs based on these models present novel strategies for identifying TCRs that bind to specific epitopes. Additionally, researchers are making innovative attempts to discover appropriate TCRs. For example, TCRPPO is a reinforcement learning-based framework to mutate an existing TCR sequence to enhance its affinity for a given epitope [29]. However, this optimization method introduces the prediction results of the classification model as the optimization index, potentially leading to instability in the model’s final performance due to the poor generalization of the classification model. ERTransformer utilizes two pre-trained BERTs to generate TCRs, in combination with a given epitope de novo [30]. Although ERTransformer has demonstrated satisfactory performance in several specific tasks, its efficacy remains to be fully verified due to the absence of comparable models for benchmarking. Aside from the models mentioned above, there have been no further significant studies on the acquisition of TCRs, which is incongruent with the crucial role TCR holds in immunity. Here, we propose GRATCR, a novel framework for de novo sequence generation, to end-to-end generate CDR3-beta of TCRs for given epitopes. Due to the crucial role of the CDR3 region in recognition and the current lack of information about alpha sequences [27, 31], our study specifically focused on the CDR3 region of the T cell receptor beta sequence. Overall, GRATCR comprised Epitope-BERT and TCR-GPT. The advances of GRATCR can be summarized in two aspects: Firstly, by integrating epitope encoder and GPT-derived TCR generator, without extra generation head, the pre-training of GRATCR only involved 1.5 million epitopes and 3 million TCRs respectively, showing its efficient use of data. Secondly, we employed a more efficient method named “Grafting” to integrate Epitope-BERT and TCR-GPT after pre-training [32]. In this study, to assess GRATCR’s performance, we employed ERTransformer as a baseline, conducted fine-tuning on the same dataset, and generated TCRs for the same epitopes. We utilized three classification models, which excel at predicting TCR-epitope binding specificity, to evaluate whether the model-generated TCRs specifically bind to the corresponding epitopes. Experimental results indicate that our model outperforms ERTransformer with the binding probability increasing by 20%. We compared the generated TCRs with natural TCRs and found that GRATCR-produced sequences exhibit superior biological function. Through sequence level and structural level analysis, we found the generated sequences covered a larger TCRs’ sequential space, while keeping high reliability. Moreover, we examined the cross-attention scores of GRATCR, reporting a remarkable concordance to crucial binding sites.

## Results

### The framework of GRATCR for end-to-end sequence generation

To enable end-to-end generation of TCR sequences for given epitopes, we propose a deep learning framework named GRATCR. Inspired by ERTransformer [30], GRATCR utilizes a hybrid strategy involving two pre-trained transformer-derived modules, as illustrated in Figure 1. However, GRATCR adapted optimization in two crucial aspects. Firstly, we pre-trained a TCR model based on the architecture of GPT rather than BERT, as decoders of GPT are inherently better for sequence generation [33]. Secondly, after re-evaluating and exploring the effects of different communication strategy between two pre-trained models, we adopted a grafting linkage strategy between Epitope-BERT and TCR-GPT. In comparison to ERTransformer that links two pre-trained encoders (Epitope-BERT and Receptor-BERT) via a randomly initialized “cross-attention” module within the Receptor-BERT, this grafted framework could maintain the robust embedding abilities of the pre-trained model [32] and generate high-quality TCRs with strong interaction capabilities toward desired epitopes. We assume that these improvements guarantee the superior performance of GRATCR despite its small pre-training scale. We have previously investigated whether the ERTransformer architecture can maintain high performance with significantly smaller pre-training scales. However, the results indicate that the quality of generated sequences deteriorates (Figure S2), suggesting that ERTransformer lacks data efficiency. Additionally, our attempt to substitute the TCR-GPT in GRATCR with a BERT model yielded unsatisfactory results as the model failed to generate sequences properly. All of these findings support the rationale behind the GRATCR framework.

**Figure 1.**
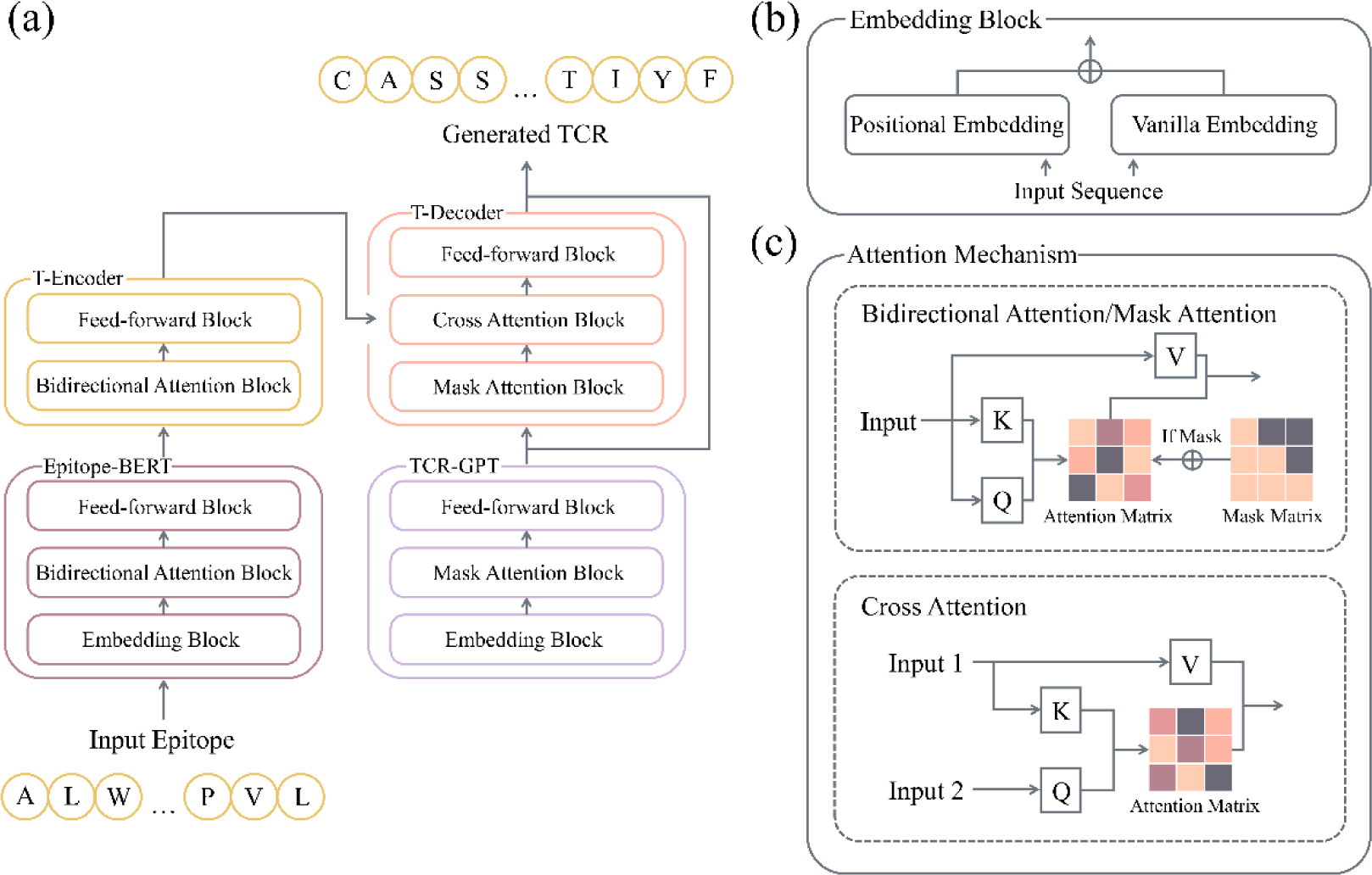
Illustrated diagram of GRATCR framework. (a) Architecture of GRATCR based on pre-trained encoder (Epitope-BERT) and decoder (TCR-GPT), connecting via a grafting mechanism. (b) The embedding block which takes sequence as input to obtain residue level representation with positional information. (c) Attention mechanism applied in GRATCR. For Epitope-BERT, bidirectional attention enables retrieval of masked token utilizing bi-directional information. In TCR-GPT module pre-trained on autoregressive task, the attention block contains mask mechanism to avoid data leakage. And the cross attention enables the communication between Epitope-BERT and TCR-GPT.

The training of GRATCR involves two stages: 1) pre-training of Epitope-BERT and TCR-GPT, and 2) SeqToSeq fine-tuning. In the pre-training stage, we utilized several public datasets comprising a large number of sequences for both epitopes and TCRs. The pre-training processes for Epitope-BERT and TCR-GPT are distinct. For Epitope-BERT, tokens from the input sequence are randomly masked with a certain probability following standard BERT’s training scheme. The model is trained to accurately recover the masked tokens, enabling it to learn the intrinsic relationships between amino acids and the biological features contributing to immunogenicity in epitopes. For TCR-GPT, we employed a ’predict the next token’ objective. This autoregressive strategy aims to minimize the discrepancy between the predicted next token and the ground truth, given the preceding tokens. This task enables TCR-GPT to learn the characteristics of TCR sequences and adapt to the sequence generation early on without the need for an additional linear generation head before fine-tuning.

We then fine-tuned the pre-trained model using a SeqToSeq task. In order to generate TCRs towards specific epitopes, the encoder and decoder should be linked to form an end-to-end framework. In ERTransformer, the two encoders are linked via an extra cross-attention module with random initialized parameters. Specifically, the output embeddings of Epitope-BERT are taken as inputof the pre-trained Receptor-BERT to provide the information of epitopes as prior knowledge. However, this strategy has been found to disrupt the decoder’s architecture during the fine-tuning stage. Such architectural perturbation can cause the model to forget valuable information related to TCR properties learned during the pre-training stage, thereby limiting its ability to generate TCR sequences effectively [32]. To address this issue, we utilized a connection strategy named “Grafting” to link Epitope-BERT and TCR-GPT. The intuition is to preserve the invariability of the pre-trained model. To achieve this, we stacked the encoder of the transformer (T-Encoder) upon Epitope-BERT and the transformer decoder (T-Decoder) upon TCR-GPT, linking these two parts together. This grafting scheme prevents direct message passing through inner-module mechanisms (i.e., TCR-GPT’s cross-attention). The fine-tuned model thus functions as an end-to-end TCR generator. To generate various TCRs that bind to specific epitopes, we employ beam search algorithms during the sequence generation process.

### TCRs generated by GRATCR exhibit higher specificity towards epitopes than the competing method ERTransformer

The primary objective of GRATCR is to generate TCRs capable of binding to specific epitopes following SeqToSeq fine-tuning. In the task of TCR recognition of epitopes presented by the antigen-MHC complex, there has been a pronounced shift in focus towards improving the accuracy of predicting TCR-epitope binding, rather than de novo generation of TCRs. Numerous deep learning models have been proposed for predicting the specific binding of T cell receptors to given epitopes [21, 24, 25, 34–38]. These models have demonstrated excellent performance in their respective tasks, providing valuable tools for assessing the quality of generated T cell receptor sequences.

First of all, we benchmarked the quality of generated TCRs from GRATCR and ERTransformer using epitopes from MIRA dataset. Due to the extremely high resource demands of wet lab evaluation, we scored the generation quality using machine learning-based discriminators. We selected ATMTCR [38], ERGO [24] and TEPCAM [25], as these discriminators are based on different architectures and have shown state-of-the-art performance in determining binding between TCRs and epitopes. We trained them on a curated MIRA dataset with negative pairs generated by shuffle, and then utilized them to evaluate the generated TCRs. The results predicted by ATMTCR showed that 71.3% of TCRs generated by GRATCR can specifically target corresponding epitopes while ERTransformer achieved 58.0% (Figure 2a). According to ERGO, the binding probability of sequences generated by ERTransformer and GRATCR are 56.6% and 66.1%, respectively (Figure 2a). The TEPCAM results showed that, on average, 52.7% of T-cell receptor sequences generated by ERTransformer could bind to corresponding epitopes, while GRATCR achieved 66.4% (Figure 2a). Furthermore, in order to ensure the objectivity of the discriminators’ scoring, we opted to re-train them using an additional merged dataset consisting of VDJdb, McPAS and IEDB, in order to mitigate inaccuracy stemming from distribution bias. The results continue to affirm the stability and superior performance of GRATCR compared to ERTransformer (Figure 2a). It is worth noting that we have observed a significant decrease in the performance of ERTransformer compared to that reported in the original paper, which may be attributed to two factors. Firstly, the performance of ERTransformer in the original paper relies on the predicted results of ER-BERT-BSP [30], which is derived from ERTransformer itself and this “homologous” setting might overestimate ERTransformer’s performance due to capturing sequence pattern biases during pre-training, necessitating other classifiers to further verify the quality of the generated sequences. Simultaneously, the introduction of exogenous TCRs and epitopes to serve as negative pairs in the training process utilized by ER-BERT-BSP is an unreliable practice. It’s proved that pairing background TCRs with exist epitopes as negative data could lead to shortcut learning [39], where the discriminator captures differences in data distribution, causing it to favor epitopes labeled as positive samples in training. To this end, we also re-trained ATMTCR, ERGO and TEPCAM with the implementation details of these discriminators are comparable to those of ER-BERT-BSP. As shown in Figure 2b, the discriminator therefore gives more attractive but deceptive results, which is a pitfall.

**Figure 2.**
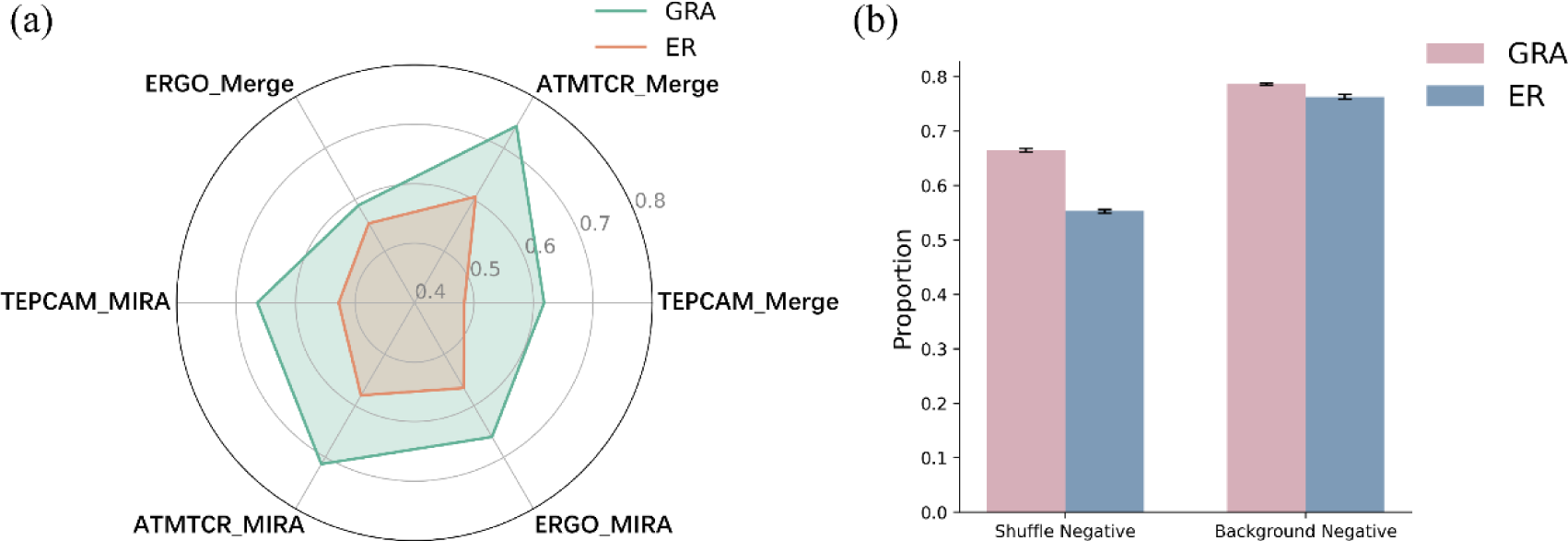
Benchmark of generated sequence quality via diverse discriminators. (a) Proportions of TCRs that could bind to specific epitopes generated by GRATCR and ERTransformer, respectively. Three models (ERGO, ATMTCR, TEPCAM) was used as external discriminators. (b) Proportions of TCRs that could bind to specific epitopes generated by GRATCR and ERTransformer on average predicted by ERGO, ATMTCR, TEPCAM trained with two negative sampling methods (shuffle, background).

In conclusion, TCRs generated by GRATCR exhibit a higher degree of specificity towards corresponding epitopes.

### TCRs generated by GRATCR exhibit higher biological conservation compared with natural sequences

The development of immunotherapies in real-world settings demands that TCRs exhibit high levels of immunogenicity and bioactivity. Specific motifs exert a crucial role in maintaining biological activity; thus, biological conformance is of paramount importance, which can be computed by BLOSUM scoring matrix value, this has been demonstrated to be efficacious in antibody design [40, 41]. To further evaluate the generated T-cell receptor sequences, we employed the BLOSUM62 scoring matrix to quantify their biological propensity compare to the natural TCRs. Higher BLOSUM62 scores indicate more biologically favorable substitutions, aiding in the identification of conserved regions and functional domains across protein sequences. As before, we used ERTransformer and GRATCR to generate T-cell receptor sequences for epitopes on the MIRA dataset and calculated their BLOSUM62 scores. For comparison, we also generated random amino acid sequences for the same epitopes as a control. Considering the relatively fixed patterns observed in the first few residues (“CASS”) and the last two residues (“YF”) of the CDR3 region of the TCR beta chain, the random sequences generation process was narrowed down to the middle part of the TCRs. Additionally, we constrained the length of the randomly generated sequences to the range from 14 to 16 amino acids, making them similar to natural sequences in terms of basic amino acid patterns. We used natural TCRs as reference sequences to calculate the average substitution score. The results showed that the average BLOSUM62 score for randomly generated sequences was 42.77 with a standard deviation of 2.70. In contrast, the sequences generated by ERTransformer had an average score of 48.67 with a standard deviation of 3.35, while those generated by GRATCR had an average score of 49.56 with a standard deviation of 3.53 (Figure 3a). This indicate that T-cell receptors produced by both ERTransformer and GRATCR are of significant biological relevance (p < 0.0001), with GRATCR-generated sequences being more favorable in evolutionary terms than those generated by ERTransformer.

**Figure 3.**
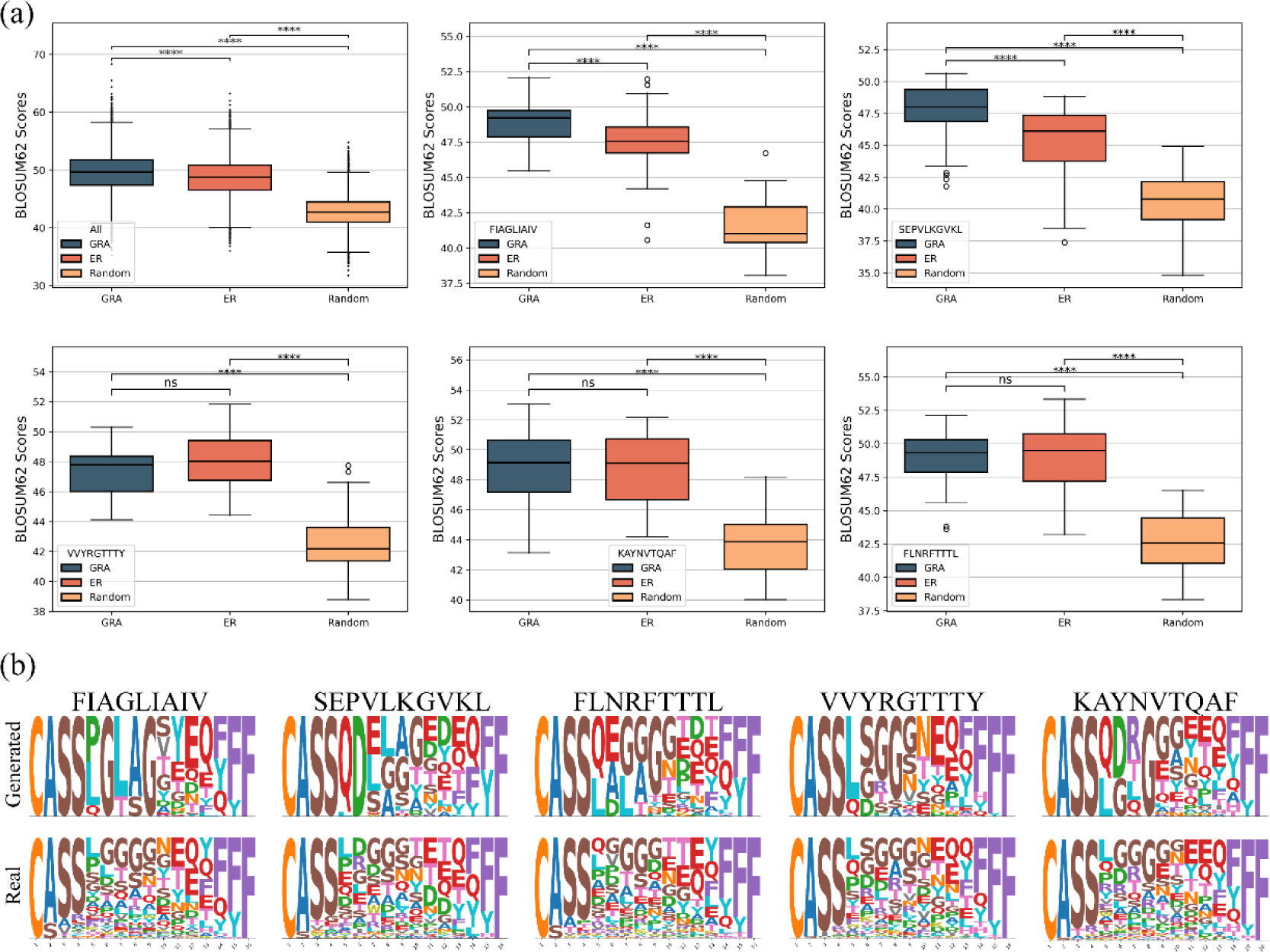
Analysis of biological and sequence level conservation of TCRs. (a) The BLOSUM62 substitution scores by comparing GRATCR generated, ERTransformer generated and random generated TCRs. ****p < 0.0001, ns.Not significant, Wilcoxon signed-rank test. (b) Amino acids composition analysis for generated and natural TCRs, in which five epitopes were picked. The height of logo position is correlated to amino acid’s frequency in particular position.

Then we analyzed the characteristics of the generated sequences, and selected five epitopes with sufficient natural TCRs paired in the dataset for specific analysis. Overall, the results for these epitopes are consistent with the population-level findings (Figure 3a). Sequences generated by GRATCR for the epitopes FIAGLIAIV and SEPVLKGVKL outperform those generated by ERTransformer. For the epitopes FLNRFTTTL, KAYNVTQAF and VVYRGTTTY, the sequences produced by GRATCR are comparable to those from ERTransformer. Regardless of the epitopes, randomly generated sequences scored significantly lower than those generated by both ERTransformer and GRATCR. Subsequently, we focused on the T-cell receptor sequences produced by GRATCR. For the five selected epitopes, we identified natural and generated T-cell receptors ranging from 14 to 16 amino acids in length, and then utilized Logomarker visualization to assess the conservation of the generated sequences (Figure 3b). In general, the generated CDR3 sequences closely resemble natural sequences. For instance, all generated sequences start with “CASS” and end with “YF” or “F,” aligning with the common pattern of T-cell receptor beta sequences. Although GRATCR appears to have captured these patterns, it’s concerning that the generated sequences were more conserved and exhibited less diversity than natural sequences. We assumed that one reason for the reduced diversity of the generated sequences might be attributed to the relatively small scale of the pre-training data, which restricts the model’s ability to learn and incorporate low-frequency amino acids. For example, in TCRs generated for the FIAGLIAIV epitope, glycine (G) was fixed in the sixth position, whereas various amino acids occupy this position in natural sequences. We analyzed the amino acid composition of the 3 million pre-training T-cell receptor sequences and discovered that glycine was the most frequent amino acid in the sixth position of TCR beta chain CDR3. Similarly, glycine frequently appears in the ninth position of sequences generated for the FIAGLIAIV epitope and in the eighth position of sequences generated for the KAYNVTQAF epitope, consistent with the high frequency of glycine in these positions in the pre-training data. These findings support our hypothesis that the pre-training data scale restricts the diversity of generated sequences. In general, while the sequences generated by GRATCR are bio-functional, they remain more conserved than natural sequences.

### TCRs generated by GRATCR exhibit novelty compared to natural T-cell receptors

Extracting connotative features of TCR holds immense significance in understanding its revolution preference and high sensitivity toward ligand epitopes [42]. Pre-training on a tremendous sequence pool helps model to learn knowledges about evolutional and functional plausibility rather than sequential patterns. The enriched prior knowledge is the basis of capacity for generator to discover a larger sequence space. Here, we extract the intrinsic characteristics of both generated and natural sequences, allowing for a comparative analysis. We firstly introduce TCR2vec [43] as a feature extractor, a pre-trained transformer-based model that encodes TCR sequences into high-dimensional vectors. Upon acquiring the sequence-level representations of both generated and natural TCRs, we employ the Adjusted Rand Index (ARI) to evaluate the similarity between the sequence clusters of natural and generated origins. ARI, a statistical measure for assessing similarity between two data clusters, ranges from -1 to 1, where a score near 1 indicates high agreement, a score near 0 suggests random clustering, and negative scores denote disagreement. We selected 20 epitopes and computed the ARI values between generated and natural sequences (Table 1). Overall, based on the characterization derived from TCR2vec, we infer that neither GRATCR-generated nor ERTransformer-generated sequences closely resemble natural ones, although for some epitopes GRATCR-generated TCRs exhibited greater similarity to native T cell receptor sequences than those generated by ERTransformer. The extensive data used in training deep models may introduce significant biases, affecting the representativeness and accuracy of feature extraction. To mitigate this influence, we then employ a sequence-based feature by specifically calculating 3-gram amino acids composition. The statistical results align with TCR2vec’s finding, indicating that generated sequences exhibit low similarity to natural sequences, which means our de novo protein sequence generator is capable of exploring a larger sequence space while maintaining sufficient biological relevance.

**Table 1.**
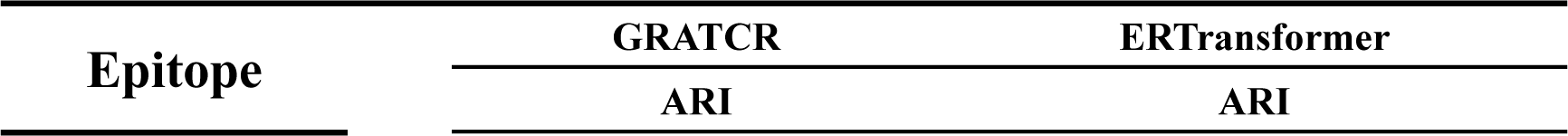
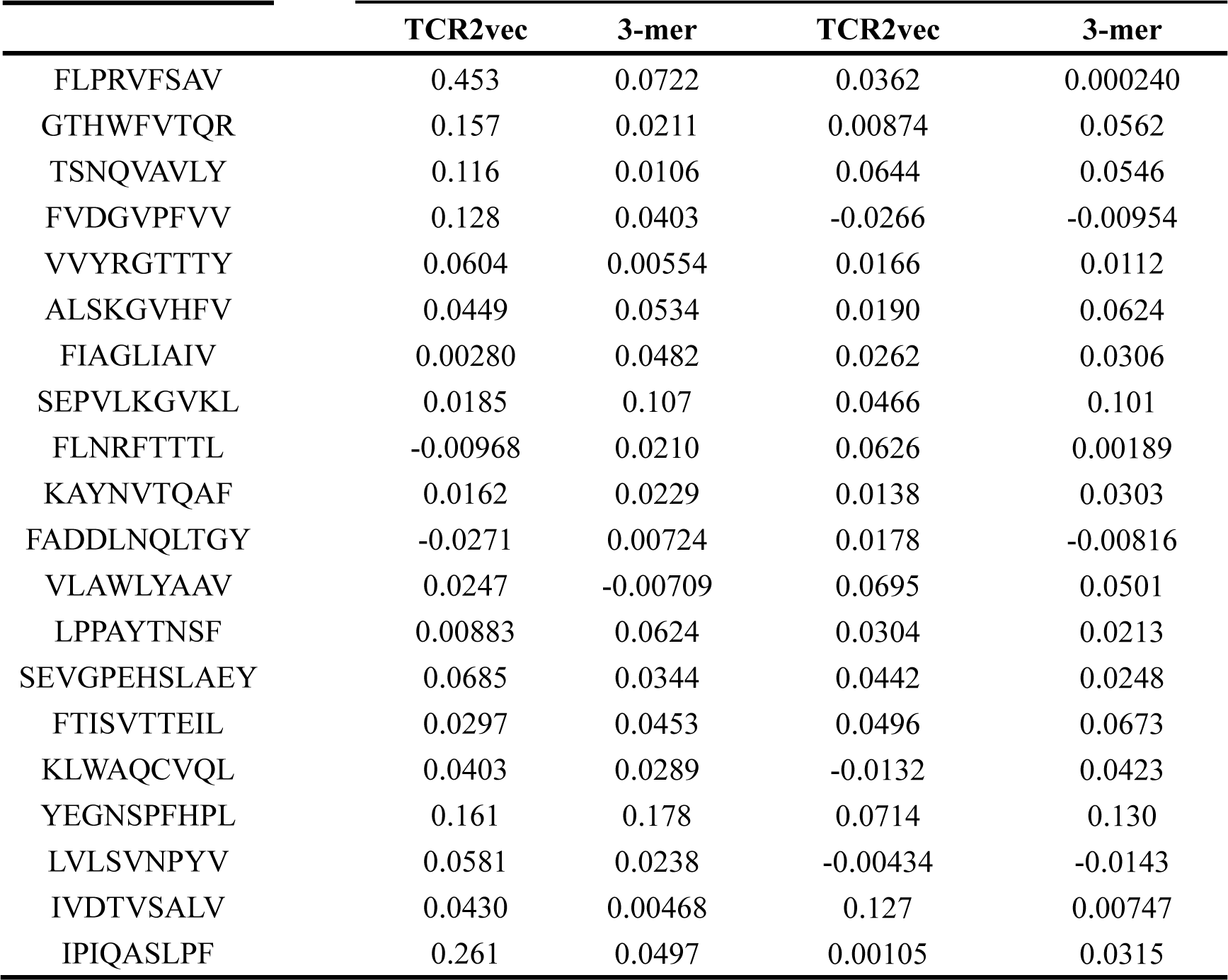
Adjusted Rand Index (ARI) values between natural TCRs and those generated TCRs using two different feature extraction methods.

### TCRs generated by GRATCR facilitate the assembly of the TCR-peptide-HLA complex

To investigate whether the generated T-cell receptor (TCR) sequences can specifically bind to given epitopes, we conducted a structural analysis of the sequences. We selected five TCR-pMHC complexes from the Protein Data Bank (PDB) with different epitopes. The epitope sequences and the associated PDB codes are ’ALWGFFPVL’ (PDB ID: 2J8U), ’EAAGIGILTV’ (PDB ID: 4QOK), ’LLFGYPVAV’ (PDB ID: 1QSF), ’LGYGFVNYI’ (PDB ID: 3PWP), and ’MVWGPDPLYV’ (PDB ID: 5C0A). For each of the five selected epitopes, we used GRATCR to generate corresponding TCRs. We then replaced the sequence of the CDR3 region in the natural TCR with the generated TCR sequence (Figure 4a). To obtain structural information about these modified sequences, we employed AlphaFold3 [44], which has achieved significant breakthroughs in predicting biomolecular structures and interactions, particularly in immunology-related sequences, such as antibody-antigen complexes, where it has demonstrated substantial improvements. To evaluate the predicted complex structures and their interactions, the Solvent Accessible Surface Area (SASA) [45] was calculated for assessing protein structure and function. First, we calculated the SASA value of the TCR-peptide-HLA complex after replacing the CDR3 region with the generated sequence (Figure 4b). Surprisingly, a significant decrease in SASA was observed compared to the natural CDR3, suggesting that the generated TCRs create a more tightly bound complex, thus ensuring the structural stability of the protein sequence after folding. Next, we focused on the interaction between the TCR sequences and the epitopes by calculating the change in SASA before and after the combination of the TCR beta region and the epitope (Figure 4c). The complex with generated CDR3 exhibited higher value than natural one, reflecting a larger interaction interface and closer binding between the generated TCRs and the epitopes. This finding aligns with the binding prediction of the discriminators in the previous exploration. Overall, the predicted results demonstrate that the generated TCR sequences help stabilize the complex structure, validating the structural integrity and binding potential of the sequences produced by GRATCR.

**Figure 4.**
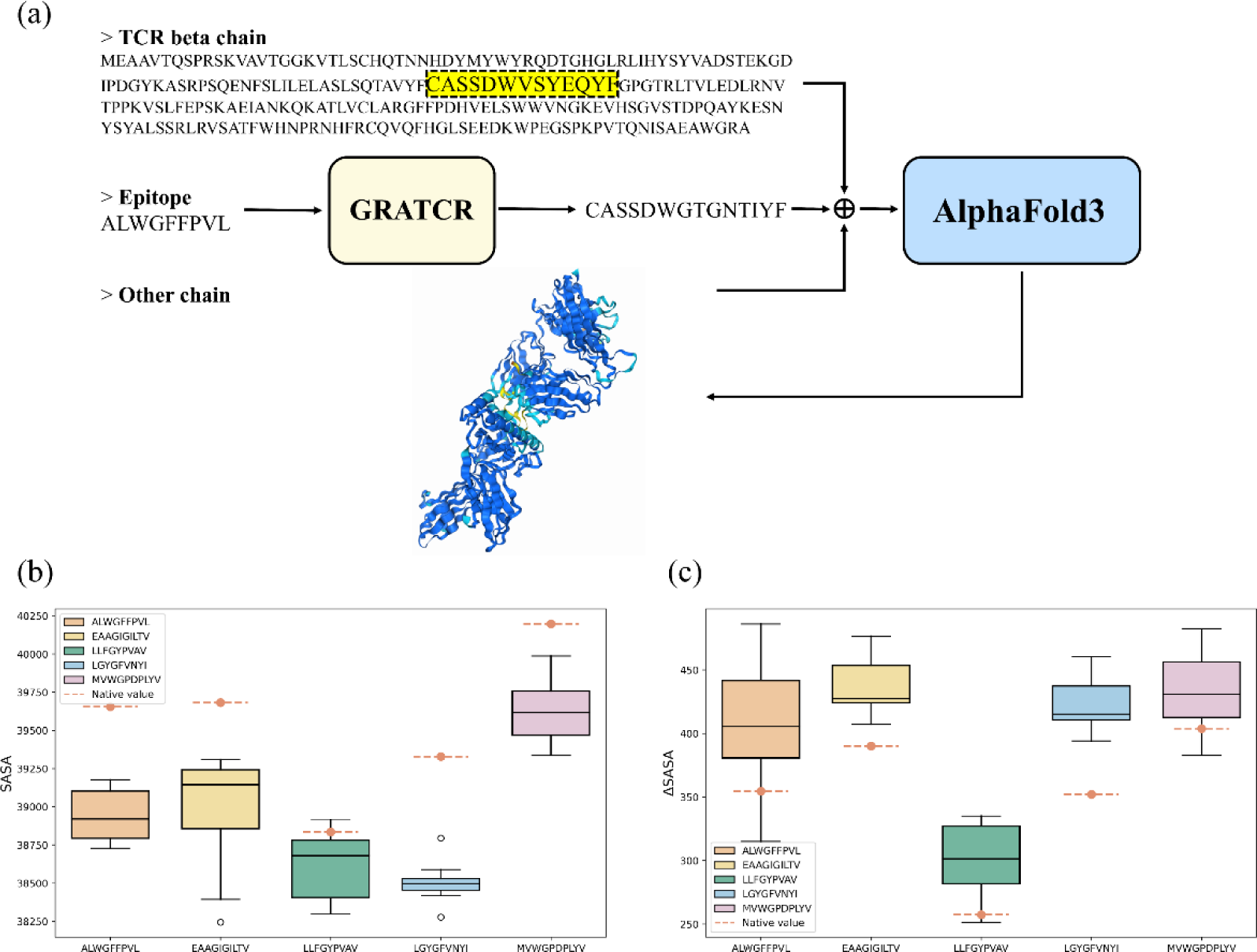
Structural analysis for quantifying binding potential. (a) The CDR3 region of beta chain in the TCR-pMHC complex was replaced by GRATCR generated CDR3. The structure of new complex is predicted utilizes the cutting-edge tool, AlphaFold3. (b) Solvent Accessible Surface Area (SASA) values of the TCR-peptide-HLA complexes with five different epitopes. (c) Change of SASA upon binding. The box plots indicate values of generated sequences, and dash line in orange is the value of original sequence.

### GRATCR can capture amino acid residues that are critical for binding

Besides the generation of TCRs with high potential to interact with given epitopes, GRATCR could provide physically meaningful insights via its explainable module. The model incorporates attention mechanism, allowing us to elucidate the knowledge acquired by the model during training. We extracted features from the cross-attention layers between the T-Encoder and T-Decoder to determine whether these features reveal biologically meaningful insights. We selected two TCR-epitope pairs, with PDB IDs 1OGA and 1LP9, as examples. Notably, neither of which were present in the training set. In the 1OGA complex, phenylalanine (PHE-7) of epitope and arginine (ARG-98) of CDR3 are in close proximity (<6Å), indicating a key site for T-cell receptor and epitope recognition. Visualization of the attention matrix shows that the sequences “SR” and “FT” have the highest attention scores (Figure 5a), suggesting that the model recognizes the importance of this interaction. Similarly, in the 1LP9 complex, phenylalanine (PHE-5, PHE-6) and tryptophan (TRP-97) are very close in 3D space, with the aromatic ring of phenylalanine (PHE-5) oriented towards tryptophan. The highest attention scores for “DW” and “FF” in the visual matrix are consistent with this observation (Figure 5b), indicating that the model also catches regions crucial for specific recognition. These findings demonstrate that our model learns some knowledge that governed mechanisms of TCR and epitope binding, enabling GRATCR to generate T-cell receptor sequences with satisfactory biological function and binding ability.

**Figure 5.**
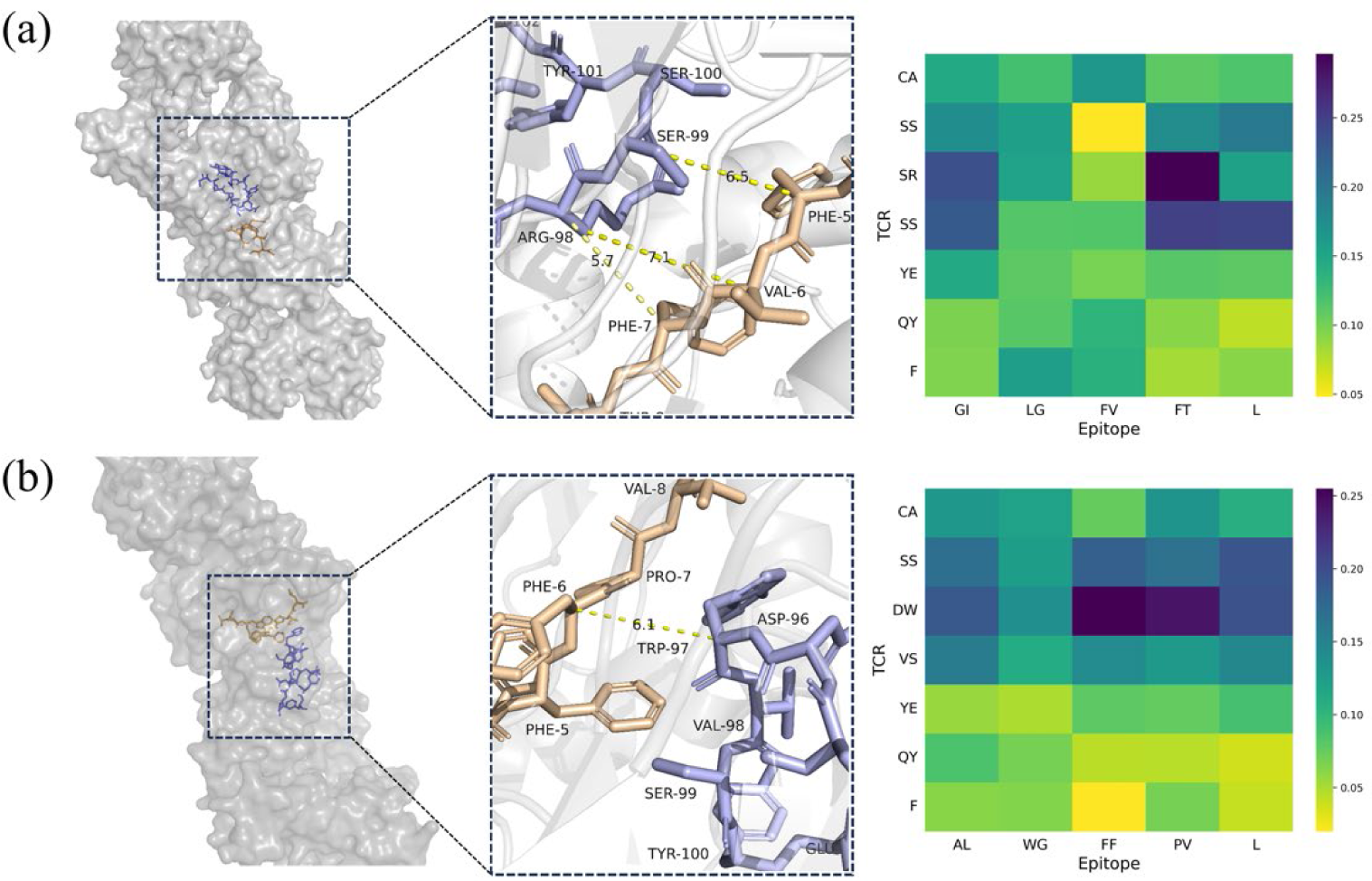
Three-dimension structure analysis and the corresponding attention scores. Left: Visualization of TCR-pMHC complex; Right: Attention scores extracted from cross-attention module between grafted T-encoder and T-decoder. (a) Complex with PDB code 1OGA, (b) Complex with PDB code 1LP9.

## Discussion

Immunology is a prominent focus in current research, and historically, numerous immune-based tumor treatments have made significant strides. Within immunology, T cell receptors are of great interest due to their pivotal role in the specific recognition of antigens. There are already therapies that modify T cells to target tumor antigens through the introduction of T cell receptor genes, yielding promising results. However, current technology still faces challenges, and the identification of TCRs capable of recognizing specific antigens heavily relies on labor-intensive wet experiments. Although there have been some attempts to apply increasingly advanced deep learning to identify TCRs for a given epitope, the lack of relevant research belies the importance of the issue.

Here, we propose GRATCR to generate TCRs de novo for a given epitope. The core components of GRATCR are Epitope-BERT and TCR-GPT, pre-trained with 1.5 million epitope and 3 million TCR, respectively. After pre-training, we attach T-Encoder and T-Decoder to the last layer of Epitope-BERT and TCR-GPT respectively to construct GRATCR. The cross-attention layer between T-Encoder and T-Decoder enables the information of epitopes to be transmitted to the decoder. In order to evaluate the performance of the model, we introduce ERTransformer as benchmark and three classifiers to predict whether the generated TCRs can be combined with corresponding epitopes. We noted that the method ER-BERT-BSP utilized for generating negative samples inclined the classifier to score higher, and the seductive results warn us against the trap of the classifier falling into the shortcut learning. We found that when generating negative samples of the MIRA training dataset, ER-BERT-BSP did not randomly shuffle the TCRs and epitopes to mismatch them, but chose to introduce sequences from a larger exogenous pool of T-cell receptors that did not bind specifically to the epitopes in MIRA to act as negative samples. At the same time, in order to increase the diversity of epitopes, many epitopes and TCRs that do not bind to each other are also introduced together as negative pairs [30]. Intuitively, this approach to increase the diversity of sequences in the training dataset seems feasible, because current classifiers have been found to be poor in generalization and highly dependent on data [46]. When the input sequences of either TCRs or epitopes are not present during training, it is difficult for model to tell whether they can bind to the corresponding epitope or TCR and we indeed found that the sequences generated by both ERTransformer and GRATCR had very few duplicates with those in MIRA. But this approach is also dangerous. It can introduce many bias that destroy the performance of classifiers. At the same time, we introduced ATMTCR, ERGO and TEPCAM and trained them with two different datasets respectively and the negative samples of the training datasets are coming from random shuffle and mismatch which is more reliable. The results indicated that TCRs generated by GRATCR are indeed more likely to combine with corresponding epitopes than those generated by ERTransformer. Although the results of the classifier show excellent performance of the model, because current prediction models are still found to be dependent on the data and have poor generalization, it is not enough to rely solely on classification predictions to evaluate whether the generated TCRs will bind specifically to a given epitope. More and more reliable studies are needed in the future to truly measure the performance and reliability of different generators. In addition, the affinity between TCR and epitope must be kept within a certain range, too high or too low affinity will lead to abnormal immune activation, only a certain range of TCR can help the body kill tumor cells without destroying normal tissue [47, 48]. However, the current public data set can only support the classification model to make zero-one prediction, and the investigation of affinity is still very lacking, so more attention and exploration are still needed. At the same time, sequence alignment was utilized to compare the generated sequences from GRATCR and ERTransformer with the natural sequences, and BLOSUM62 scores were calculated to evaluate the bio-functionality of the sequences. In order to increase the control, we also randomly generate some sequences, which mimic the pattern of TCR. The final results show that the functionality of our GRATCR-generated sequences is better than that of the other two groups. In order to specifically analyze the characteristics of the generated sequences, we selected some epitopes to explore that, by calculating the relative frequency of amino acids occurring at each position, the generated sequences were indeed lower in abundance than the natural sequences. Simultaneously, we employed two methods to extract feature vectors of the sequences to compare the similarity between natural and generated sequences. The results indicate that the generated sequences are novel. Additionally, we analyzed the structural aspects of the generated sequences using AlphaFold3. The results demonstrated that the generated sequences could stabilize the structure of the complex and increase the interaction area, further supporting the quality of the generated sequences. At the same time, we also analyzed the attention mechanism inside the model, and the model can indeed capture some decisive sites, which supports the high score of the classifier is trustworthy to a certain extent. However, as mentioned earlier, the evaluation of the generator’s performance requires more exploration, and the current conditions still limit our in-depth analysis. Overall, our work adds some new ideas to related tasks (i.e., the generation of new TCRs), which is beneficial for T-cell based therapy.

## Materials and methods

### Datasets

To pre-train Epitope-BERT and TCR-GPT, we selected two datasets: IEDB [49] and TCRdb [50]. The Immune Epitope Database (IEDB) serves as an extensive repository of immune epitopes, from which we curated 1.5 million epitopes to pre-train Epitope-BERT. The TCRdb encompasses a substantial collection of unlabeled TCR sequences obtained via TCR-seq technology. We curated 3 million TCRs from this database to pre-train the TCR-GPT module. Following pre-training, we utilized the MIRA [51] dataset to fine-tune the model. The MIRA dataset contains epitopes related to the SARS-CoV-2 virus and the corresponding T cell receptors. From this dataset, we retained 151 epitopes along with 43,983 TCR-epitope pairs to fine-tune GRATCR. In the subsequent generation process, we generated T cell receptor sequences for the 151 epitopes in the MIRA dataset. Additionally, to train the external discriminators, we merge a dataset consisting of VDJdb[52], McPAS[53] and IEDB.

### Overview of GRATCR

Herein, we introduce the framework of the model in detail. The key components of GRATCR are Epitope-BERT and TCR-GPT that are designed to catch the rules of epitopes and TCRs respectively. Each of them is pre-trained with different tasks. In pre-training, when given an epitope sequence, some tokens in the sequence are randomly masked and entered into Epitope-BERT’s embedding layer, which includes both vanilla and positional embedding. The sequence then enters the Epitope-BERT blocks with 8 layers of standard BERT blocks. Each block contains a multi-head attention layer and a fully connected feedforward network, in which the multi-head attention mechanism is the core part of the overall framework, helping the model to grasp the information contained by the individual amino acids in the sequence and their relationships to each other. The attention score of a single head can be calculated using formula (1), where Q represents Query, K represents Key, V represents Value and d_k_ represents the dimensionality of the Key.

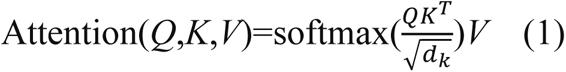

The final output result of the multi-head attention layer can be expressed by formula (2), and the attention score of each head is calculated as shown in formula (3) where 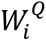, 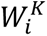 and 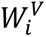 are the learned linear transformations for each head.

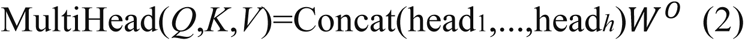

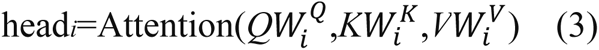

After calculating the sequence vector through all blocks, we use cross entropy as a loss function to calculate the loss between the masked tokens and the output of the model to update the weight. When pre-training TCR-GPT, the input TCR sequence also passes through the same embedding layer including vanilla and positional embedding. The processed sequence vectors then pass through eight standard GPT blocks, each containing a masked multi-head attention layer and a fully connected feedforward network.

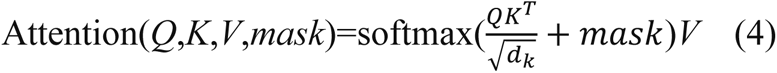

The mask attention score can be calculated using formula (4) where mask is a matrix with negative infinity values at position should be masked and it can help mask the influence of future positions in the input sequence to the front which is more in line with the situation when the sequence is actually generated. Finally, when the sequence vector has passed through all the blocks, we use cross entropy to calculate the loss between the output value and the label to update the weights.

After pre-train, we select the Encoder of transformer(T-Encoder) without embedding layer and attach it to the upper layer of our pre-trained Epitope-BERT to further process the representations of epitopes. Meanwhile, we attach the Decoder of transformer(T-Decoder) to the upper layer of pre-trained TCR-GPT. At the beginning of the finetuning, the weights of T-Encoder and T-Decoder are randomly initialized. Both T-Encoder and T-Decoder have 6 same blocks and the Cross-Attention layers between them naturally connect the whole Encoder composed of pre-trained Epitope-BERT and T-Encoder and whole Decoder composed of pre-trained TCR-GPT and T-Decoder. When finetuning, Epitope-BERT and TCR-GPT take epitopes and TCRs as input respectively and output the high-dimensional representations. Then, they are fed into T-Encoder and T-Decoder and the information of epitopes can be transferred to Decoder by Cross-Attention layers. Finally, we employ a residual connection between the final layer of TCR-GPT and T-Decoder to get final outputs.

Here, we trained GRATCR on two GeForce RTX 3080 GPUs with 20GB memory using the PyTorch 1.10.1 framework and Python 3.9 and the communication between different GPUs is realized by accelerate library. The model weights were updated by AdamW optimizer with warmup step proportion set to 0.1. Specifically, the hyperparameters of GRATCR are: learning rate = 5e-5, number of heads of Attention mechanism = 12, dimensions of embedding layers = 768, all of them are widely utilized in various applications.

### Methods of sequence evaluation

To evaluate the biological functional similarity between the generated T cell receptor sequences and the natural sequences, we calculated BLOSUM62 values. We use BioPython library’s pairwise2 module to align generated TCR sequences against a set of natural TCR sequences binding to the same epitope. The alignment scoring was based on the BLOSUM62 substitution matrix. At the same time, to evaluate the conservatism of the sequence, we utilized Logomarker, a Python library for generating sequence logos, to visualize the amino acid sequence conservation and variation after calculating the relative frequencies of amino acids at each position.

In order to assess the similarity between the generated T cell receptor sequences and the natural sequences, we employed two distinct methods to extract feature information from the sequences. Firstly, we introduced TCR2vec [43], a deep learning-based transformer-derived model that has been pre-trained for characterizing TCR sequences. Upon receiving the sequence input, TCR2vec produces a 120-dimensional sequence feature vector. Subsequently, Kmeans clustering and the Adjusted Rand Index (ARI) are applied to compare the similarity of the embeddings derived from the generated and natural sequences. The implementation of Kmeans clustering and the calculation of ARI are based on the Python library scikit-learn. Simultaneously, given the extensive use of data in pre-training for deep models, it is important to acknowledge that these data may contain inherent biases, resulting in a potential decrease in the model’s ability to accurately represent sequences. Therefore, we have also incorporated n-gram as an alternative method for characterizing sequences, particularly beneficial for processing bioinformatics data, and specific text analysis tasks due to its capability of capturing local pattern and structure information within text. In this approach, we set n as 3 and generate the feature vector by segmenting the sequence into consecutive sets of three characters. The implementation of this method is based on the CountVectorizer from the scikit-learn Python library.

To assess the structural reliability of generated sequences, we selected certain epitopes from PDB and produced T cell receptor sequences accordingly. Subsequently, we replaced the CDR3 region of natural T-cell receptor beta sequence with the generated sequence and used Alphafold3 [44] to obtain structural information. To quantify this information structurally, we visualized Alphafold3’s predicted structure using PyMol while simultaneously calculating Solvent Accessible Surface Area value - a crucial reference point for evaluating protein-protein interactions and complex formation - setting probe radius at 1.4 Å and point density at 4.

### Baseline models

To evaluate GRATCR’s performance, we employed ERTransformer[30] as a benchmark, given its pioneering contributions to the generation of T cell receptors. ERTransformer’s framework incorporates EpitopeBERT and ReceptorBERT, pre-trained on 1.9 million epitopes and 33 million TCRs, respectively. The vast scale of this dataset underpins its exceptional performance, thus providing a robust benchmark for assessing GRATCR. To verify the binding affinity of the generated T cell receptor to the epitope, we introduced ATMTCR [38], ERGO [24], TEPCAM [25], all of which are deep models that perform well in the task of predicting whether a given epitope and T cell receptor will bind. ERGO uses two different architectures to deal with TCR embedding, namely LSTM and Autoencoder. In our study, we used ERGO-AE. Both ATMTCR and TEPCAM are deep models based on attention mechanisms. In addition, to obtain the representations of T cell receptors, we introduced TCR2vec[43], a pre-trained transformer-based model that can embedding TCRs into numerical vectors.

## Author contributions

Y.X. conceived and supervised the study. Z.Z, J.C and S.L design the model and algorithms. Z.Z and J.C performed experiments, Z.Z analyzed the results and write the original draft. All authors contributed to writing and reviewed the manuscript.

## Data Availability Statement

The codes and datasets are open-source and available at https://github.com/zhzhou23/GRATCR.

## Supporting Information

Additional supporting information can be found online in the Supporting Information section at the end of this article.

