## Supplementary File for "GRATCR: epitope-specific T cell receptor sequence generation with data-efficient pre-trained models"

In the supplementary material, we have included a github repository containing the models introduced in this study and relevant references in **Table S1**. The amino acids occurring at three positions within natural sequences of lengths between 14 and 16, along with their frequencies, are presented in **Table S2**. Additionally, original SASA data is provided in **Table S3-S4**. We present in **Figure S1** the distribution of scores predicted by three classifiers trained using sequences from larger exogenous pools as negative samples. Additionally, we utilize the same pre-training scale as GRATCR to re-pre-train ERTransformer, and the length distribution of the generated sequence is depicted in **Figure S2**. We demonstrate the diversity between GRATCR-generated TCRs and natural ones in **Figure S3**, with TCRs varying in length from 14 to 16.

**Table S1. The details of** baseline models and their github repositories.

| Model | Method | Web site |
| --- | --- | --- |
| ER-Transformer[1] | Pre-train, BERT, Transformer | https://github.com/TencentAILabHealthcare/ER-BERT |
| ATMTCR[2] | Self-Attention | https://github.com/Lee-CBG/ATM-TCR |
| ERGO[3] | Pre-train, LSTM, Auto-Encoder | https://github.com/louzounlab/ERGO |
| TEPCAM[4] | Self-Attention, Cross-Attention, Multi-channel CNN | https://github.com/Chenjw99/TEPCAM |
| TCR2vec[5] | Pre-train, Transformer | https://github.com/jiangdada1221/TCR2vec |

**Table S2.** The frequency of amino acids at three locations in the pre-training data. Length of TCRs ranges from 14 to 16.

| Amino acid | Frequency of each position | | |
| --- | --- | --- | --- |
|  | 6 | 8 | 9 |
| Alanine (A) | 138320 | 169841 | 159303 |
| Arginine (R) | 189098 | 167644 | 133445 |
| Asparagine (N) | 32247 | 34119 | 124377 |
| Asparticacid (D) | 133443 | 73801 | 70781 |
| Cysteine (C) | 660 | 599 | 780 |
| Glutamine (Q) | 82931 | 87317 | 64285 |
| Glutamicacid (E) | 86772 | 58226 | 55837 |
| Glycine (G) | 382672 | 590922 | 451693 |
| Histidine (H) | 18018 | 15462 | 21947 |
| Isoleucine (I) | 32726 | 31138 | 37267 |
| Leucine (L) | 122634 | 94643 | 87982 |
| Lysine (K) | 24247 | 18603 | 28647 |
| Methionine (M) | 17323 | 13696 | 16977 |
| Phenylalanine (F) | 30592 | 24431 | 31747 |
| Proline (P) | 123916 | 65455 | 87837 |
| Serine (S) | 178190 | 180966 | 206014 |
| Threonine (T) | 149330 | 146799 | 145380 |
| Tryptophan (W) | 35897 | 18031 | 18967 |
| Tyrosine (Y) | 28719 | 23301 | 62972 |
| Valine (V) | 81335 | 73996 | 82832 |

**Table S3.** The native value of Solvent Accessible Surface Area (SASA) of five picked TCR-peptide-HLA complexes from PDB and △SASA that reflects the size of the contact area.

| Epitope | SASA | △SASA |
| --- | --- | --- |
| ALWGFFPVL | 39654.734 Angstroms^2 | 354.57 Angstroms^2 |
| EAAGIGILTV | 39681.961 Angstroms^2 | 390.14 Angstroms^2 |
| LLFGYPVAV | 38834.633 Angstroms^2 | 257.465 Angstroms^2 |
| LGYGFVNYI | 39329.156 Angstroms^2 | 352.183 Angstroms^2 |
| MVWGPDPLYV | 40197.332 Angstroms^2 | 403.846 Angstroms^2 |

**Table S4.** SASA and △SASA of TCR-peptide-HLA complexes modified with generated CDR3 beta sequences.

| Epitope | SASA | △SASA |
| --- | --- | --- |
| ALWGFFPVL | 38943.7754±173.579 | 405.8112±53.331 |
| EAAGIGILTV | 38982.8445±381.192 | 439.1005±23.582 |
| LLFGYPVAV | 38616.7168±215.170 | 300.7460±30.631 |
| LGYGFVNYI | 38504.3383±132.380 | 422.7356±20.952 |
| MVWGPDPLYV | 39611.4734±209.533 | 433.0024±33.727 |

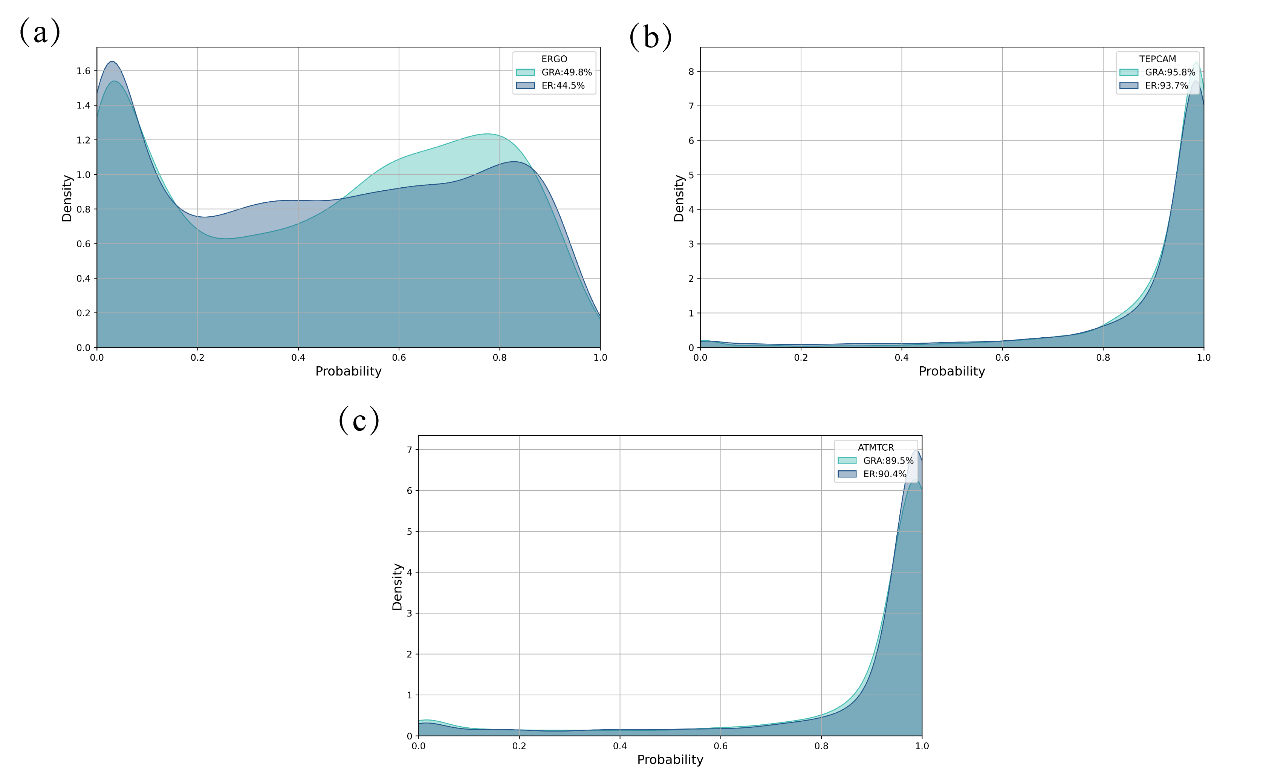

**Figure S1.** The distribution of predicted score of three discriminators trained on dataset whose negative samples are generated by introducing sequence from a larger exogenous pool of TCRs and epitopes. (a) ERGO (b) TEPCAM (c) ATMTCR.

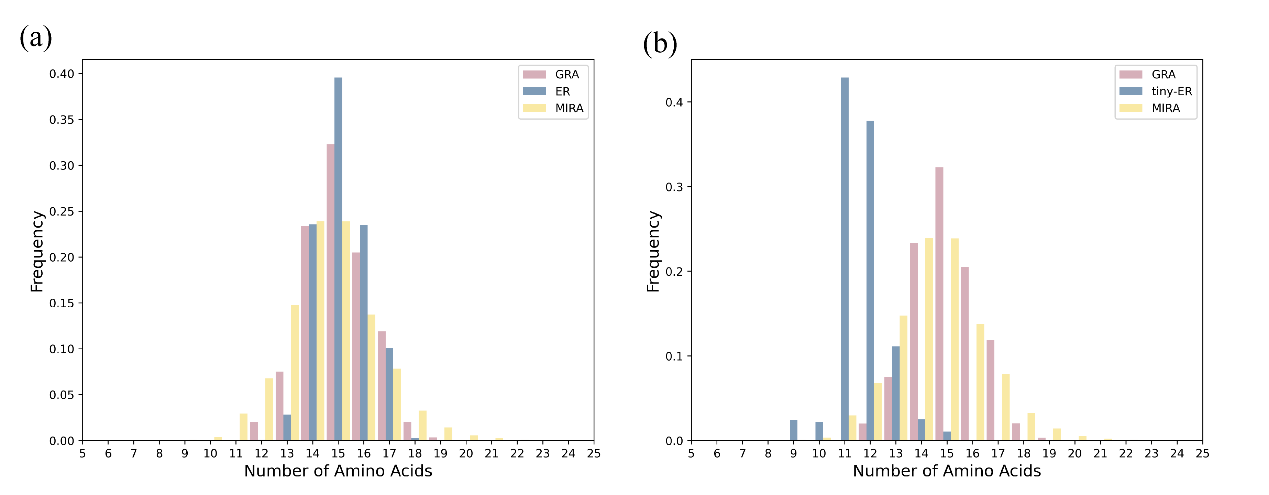

**Figure S2.** The distribution of length of TCRs from two generators (a, GARTCR and ERTransformer b, GRATCR and tiny-ERTransformer) and MIRA dataset, here tiny-ER stands for ERTransformer with pre-trained scale reduced to the same size as GRATCR.

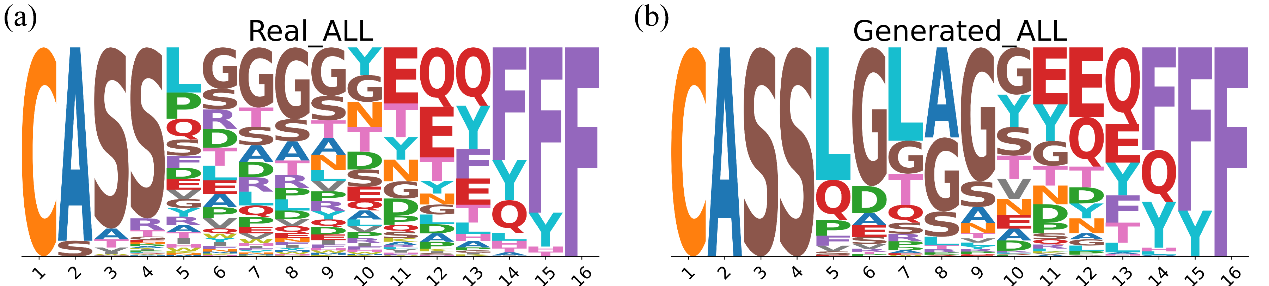

**Figure S3.** Amino acids composition analysis for all generated (b) and natural (a) TCRs, length of which ranges from 14 to 16. The height of logo position is correlated to amino acid’s frequency in particular position.
